# Discovering motifs and genomic patterns with SMT: a high-performance data structure for counting kmers

**DOI:** 10.1101/2023.04.01.535163

**Authors:** Jader M. Caldonazzo Garbelini, Danilo Sipoli Sanches, Aurora Trinidad Ramirez Pozo

**Affiliations:** Department of Informatics, Federal University of Paraná, 1299, XV de Novembro street, 80060-000, Paraná, Brazil; Department of Informatics, Federal University of Technology, 1640, Alberto Carazzai avenue, 86300-000, Paraná, Brazil

**Keywords:** kmers, motifs, ChIP-seq, sequence analysis

## Abstract

**Motivation:** The search for conserved motifs in DNA sequences is an important problem in bioinformatics. The growing availability of large-scale genomic data poses significant challenges for computational biology, particularly in terms of efficiency in analysis, kmer identification, and noise presence. The detection of conserved motifs and patterns in DNA sequences is crucial for understanding gene functions and regulations. Therefore, it is essential to develop a data structure that can handle these large volumes of information and provide accurate and fast results.

**Results:** We present SMT, an innovative tool designed to efficiently store and count kmers, optimizing memory usage and computation time. The application of SMT has also proven effective in discovering motifs in noisy datasets, allowing the identification of conserved regions in sequences. Furthermore, SMT enables exact searches in constant time and recovers the most abundant k-mers, as well as performs approximate searches in linear time to find fragments with up to *d* mutations. This approach facilitates large-scale data analysis and provides important insights into the conserved properties of biological sequences. The application of SMT in motif discovery demonstrates its potential to drive research in bioinformatics and genomics. Supplementary data and results are available to provide additional information and support the conclusions presented in this work.

**Availability and implementation:** The source code of the presented method is publicly available at https://github.com/jadermcg/SMT.

## Introduction

In recent years, there has been an exponential increase in the amount of available genomic data, thanks to advances in DNA and RNA sequencing technologies. This massive accumulation of data presents new challenges for bioinformatics, particularly in terms of efficiency in analyzing and identifying kmers and subsequences. The detection of motifs and conserved patterns in sequences is crucial for understanding gene functions and regulations, as well as for identifying functional and structural elements of the genome [Goodwin et al., 2016].

To date, various data structures and algorithms have been proposed to deal with the increasing demand for efficient large-scale kmer analysis. However, many of these solutions are not sufficiently fast or require significant computational resources, which limits their applicability to ever-growing genomic datasets [Deorowicz et al., 2019, Marchet et al., 2019].

In this context, we introduce the Sparse Motif Tree (SMT), an innovative tool specifically designed to store and count kmers efficiently. The SMT optimizes memory usage and computation time, allowing for the rapid and accurate analysis of large volumes of genomic data.

The SMT provides advanced features, such as exact search in constant time, retrieval of the most abundant kmers, and approximate search in linear time to find fragments with up to *d* mutations uniformly distributed across their bases. These features enable researchers to identify recurring patterns and conserved regions within sequences, as well as analyze variations within DNA and RNA sequences.

Therefore, the SMT is particularly useful in the problem of motif discovery, a central challenge in bioinformatics and genomics [Bailey et al., 2015]. Detecting conserved sequences and recurring patterns in DNA and RNA sequences is crucial for identifying functional elements and understanding gene regulation in different organisms. The efficiency and versatility of the SMT allow researchers to quickly analyze large genomic datasets and accurately identify biologically relevant regions with precision and reliability.

Furthermore, there is a need to deal with noisy data [Howe et al., 2021] and the SMT was designed to tackle this challenge. The ability to efficiently work with noisy data makes the SMT a useful tool for analyzing conserved regions.

The results obtained suggest that SMT and the developed algorithms have great potential to boost the search for conserved motifs and facilitate the analysis of ChIP-seq data. This work contributes to the development of new tools and methods to deal with the challenge of finding conserved regions and opens up new perspectives for the analysis of molecular biology data.

In addition to its application in motif discovery, SMT can also be used in various other contexts within bioinformatics and genomics, such as comparative genome analysis, identification of functional elements in sequences, study of genetic variations, and metagenomics [Wood et al., 2019, Chaisson et al., 2019]. The versatility and efficiency of SMT make it a valuable tool for researchers seeking scalable and high-performance solutions for handling large-scale genomic data analysis.

In this paper, we described the architecture and implementation of SMT, as well as a practical application in motif discovery. In addition, we discussed the potential of SMT to boost research in bioinformatics and genomics, helping researchers to explore and understand the complexity and diversity of genomes of different organisms.

## Implementation

SMT is a C++ application that consists of a two-dimensional data structure *M*^*ν*×6^, where *ν* represents the number of nodes, and is built from fixed-width text segments. Unlike KTrees or STrees, SMT was specifically designed to handle kmers. Its design was loosely inspired by the theory of Room squares [Archbold and Johnson, 1958], in which each element *M*_i, j_ is either empty or has a value assigned between 1 and *ν*. The main goal of SMT is to efficiently store all the kmers belonging to the main data set, as well as their addresses and counts. In this regard, each row of *M* represents a node 1,2,3,…,*ν*, columns 1 to 4 represent nucleotides, and the last two columns, 5 and 6, indicate the occurrence frequency of a segment in the data set and its respective address.

For example, if *M*_1,*T*_ = 10, the symbol *T* is present between nodes 1 and 10. Compared to other similar structures, the SMT is more efficient in storage and search of fixed-size segments, and its construction has a linear time complexity of *O*(*k*), where *k* is the size of the sequence of interest. In addition, the SMT allows for the execution of various algorithms in constant and linear time, such as exact search and approximate search, respectively, which enables the rapid retrieval of the most abundant kmers in the dataset.

The worst-case space complexity of SMT is *O*(*σν*), where *σ* is the number of symbols in the alphabet (in this case, 4) and *ν* is the number of nodes in the tree. The value of *ν* is directly dependent on the size and variance of the fragments. Thus, the greater the variance, the greater the number of nodes. It is important to note that due to its architecture, most elements of *M* will be empty. This behavior is recurrent and can be calculated. In particular, we can calculate the expected value of the occupancy level of *M*. For this, consider that the maximum and minimum number of children that a node can have is 4 and 0, respectively. The average would then be 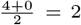 children per node.

A tree with depth k and each node averaging 2 children will have an average of 2^*k*^ leaves. The average number of nodes in the tree will be 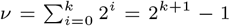. In this case, we will need a matrix *M* with 2^*k*+1^ – 1 rows and 4 columns. However, each row has 4 columns, and on average, only 2 of them will be occupied. Thus, we can calculate the occupancy level through 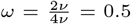. In practice, this number will be even lower, as the average number of children decreases as the depth of the tree increases. Therefore, the average space complexity of *M* is *O*(*ων*), and since *ω* < *σ*, *O*(*ων*) < *O*(*σν*).

Thus, we can expect that only half of the capacity of *M* will be used, which characterizes a significant waste of space. For this reason, SMT was implemented using a high-performance sparse matrix through the Armadillo linear algebra library [Sanderson and Curtin, 2016] using the C++ programming language. The time complexity of SMT is easier to calculate and in all cases is *O*(*mk*), where m represents the quantity and *k* the width of the fragments. To see this, we just need to consider that each fragment is composed of *k* symbols. Since there are *m* fragments, we will need to process a total of *mk* nucleotides.

One advantage of SMT is its speed compared to the construction of KTrees or STrees, whose complexity is *O*(*m*^2^) in all cases. In addition, SMT only stores the fragments of interest, making it more efficient for handling kmers. Compared to hash tables, SMT also consumes less space. Suppose we have *m* kmers of size *k* and use a direct-address hash table whose keys are the kmers. If the background distribution is uniform, the amount of space required by the hash table will be on the order of *m* × *k* bytes. In contrast, SMT only stores the necessary information and consumes less space.

## Results and discussion

In this section, we demonstrated the performance of SMT on five ChIP-seq datasets. For each dataset, we constructed the corresponding SMT, calculated its occupancy level, and reported the amount of space occupied by each one. In addition, we conducted two types of experiments: with and without noise, to demonstrate the ability of SMT to efficiently handle noisy data.

For the noise-free data, the files in the Browser Extensible Data (BED) format were obtained from the JASPAR website [Castro-Mondragon et al., 2022]. This file contains information on the genomic positions of the regions of interest. Next, we used specialized tools to extract the sequences corresponding to these regions from the human genome. Each sequence has 100 bp, and the peak regions were kept at the center.

For the noisy data, we generated *n* additional sequences based on a fourth-order Markov model whose parameters were estimated from the main dataset. These noisy sequences were then merged with the original dataset, resulting in an expanded set containing a total of 2*n* sequences. Finally, we randomly selected a sample of n sequences from this expanded set to form the noisy dataset. Thus, the noisy dataset is a combination of original sequences and sequences generated based on the Markov model, reflecting the presence of noise in the data. Table 1 shows a summary of the experiments.

**Table 1.** Comparison of SMT performance in different genomic datasets: Analysis of execution time, occupancy level, and space usage for k-mers of various sizes as reported in the literature.

| Dataset | Number of sequences | Value of k | Execution time (s) | Occupancy level | Space occupied (MB) |
| --- | --- | --- | --- | --- | --- |
| MA1637.1 | 18.823 | 10 | 12.05 | 0.0182 | 24.3 |
| MA0018.4 | 55.086 | 10 | 38.34 | 0.0081 | 32.6 |
| MA0143.4 | 62.667 | 7 | 15.36 | 0.0001 | 0.6 |
| MA0047.3 | 189.473 | 9 | 128.99 | 0.0023 | 36.4 |
| MA0148.4 | 221.646 | 10 | 143.45 | 0.0023 | 36.2 |

The tests were performed on an Intel Core i7 computer with 4 processing cores and 8GB of RAM.
The choice of different values of $k$ for each dataset was based on data collected from the literature.
The execution time was calculated by taking the average of 30 runs.

Based on the data presented in Table 1, we can make some observations and analyses about the performance of SMT on different datasets. Firstly, the execution time seems to increase as the number of sequences in the dataset increases. For example, the execution time for the MA1637.1 dataset is 12.05 seconds, while for the MA0148.4 dataset, it is 143.453 seconds. This indicates that SMT may have a performance proportional to the size of the dataset analyzed.

The occupancy level varies among the different datasets and values of k. However, in general, the values are relatively low, indicating that SMT is efficient in its use of storage space. For example, the occupancy level for the MA0143.4 dataset is only 0.0001.

The space occupied by SMT also varies depending on the dataset and the value of k. In general, the space occupied seems to increase with the number of sequences and the size of the k-mers. However, SMT still exhibits good efficiency in its use of storage space. For example, for the MA0143.4 dataset with k = 7, only 0.6 Mb of space is occupied.

Although the analysis of the algorithms involved in SMT was carried out using big-O notation, we also chose to measure the execution time empirically. This complementary approach aims to facilitate understanding and provide a more tangible perspective on the performance of SMT in practice, allowing for a more concrete evaluation of the results obtained. Table 2 shows the results of the analyses on the noisy data.

**Table 2.** Comparison of SMT performance on different noisy genomic datasets: Analysis of runtime, occupancy level, and space usage for various k-mer sizes as reported in the literature.

| Dataset | Number of sequences | Value of k | Execution time (s) | Occupancy level | Occupied space (MB) |
| --- | --- | --- | --- | --- | --- |
| MA1637.1 | 18.823 | 10 | 11.69 | 0.0182 | 30.6 |
| MA0018.4 | 55.086 | 10 | 40.21 | 0.0102 | 38.9 |
| MA0143.4 | 62.667 | 7 | 14.21 | 0.0001 | 0.6 |
| MA0047.3 | 189.473 | 9 | 102.23 | 0.0008 | 10.4 |
| MA0148.4 | 221.646 | 10 | 173.99 | 0.0026 | 40.0 |

The tests were run on an Intel Core i7 computer with 4 processing cores and 8GB of RAM.
The choice of different values of $k$ for each dataset was based on data collected from the literature.
The execution time was calculated as the average of 30 runs.

Table 2 shows the performance of SMT on different noisy genomic datasets. Comparing the results of this table with Table 1, we can observe some differences. In general, the execution time of SMT on datasets with noise is slightly higher than on datasets without noise. This can be expected since noisy data tends to have less repetition and, therefore, takes more time to analyze and construct the data structure.

The occupancy level in datasets with noise is, in some cases, slightly higher than in datasets without noise (e.g., MA0018.4 and MA0148.4). This may be due to the presence of additional k-mers generated by the noise, which increase the density of the data structure. The space occupied by SMT in datasets with noise is also generally higher compared to datasets without noise. This indicates that SMT requires more storage space to handle the additional k-mers generated by the noise.

However, despite these differences, SMT still exhibits efficient performance even in datasets with noise, demonstrating its ability to handle more complex and realistic scenarios in genomic sequence analysis. The comparison between the tables reinforces the robustness and applicability of SMT in environments with different levels of noise. Figures 1 and 2 present a comparison between the logos identified by SMT and the original logos obtained from JASPAR.

**Fig. 1.**
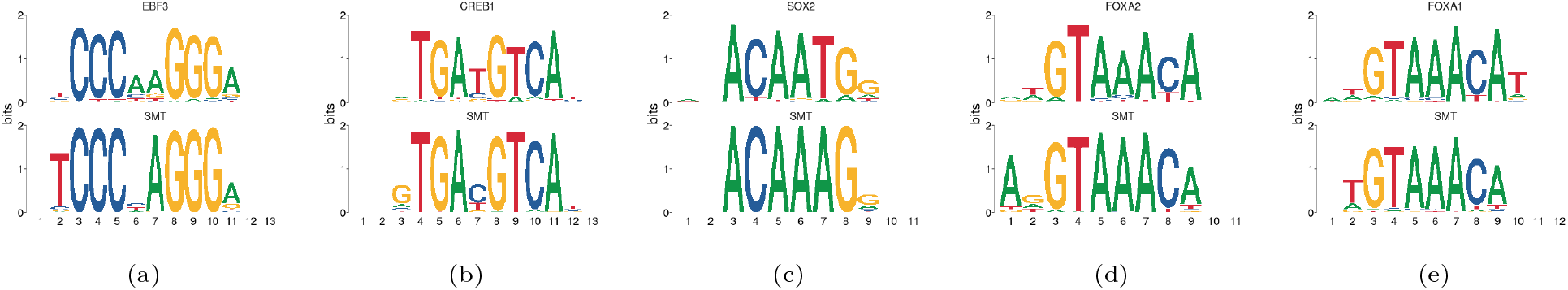
Comparison between the logos of sequences generated by SMT (below) and the logos extracted from the JASPAR database (above) for the noise-free data.

**Fig. 2.**
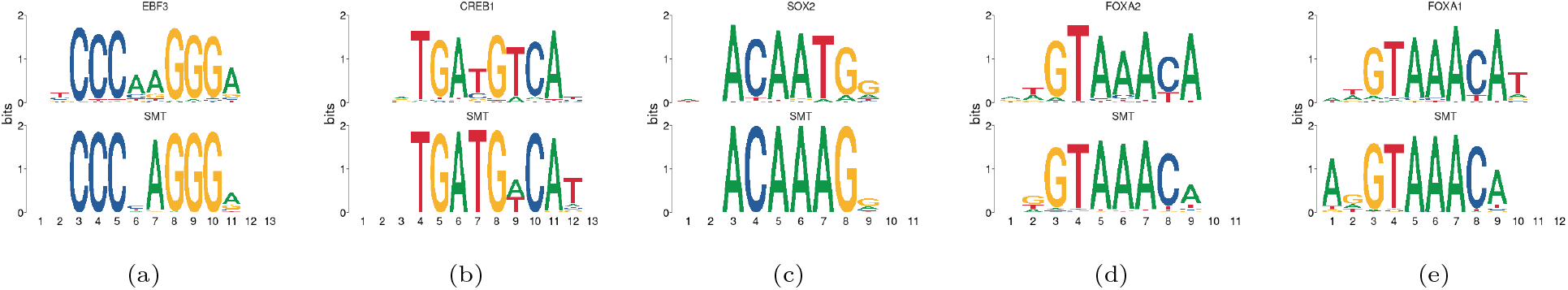
Comparison between the logos of sequences generated by SMT (below) and the logos extracted from the JASPAR database (above) for the noisy data.

The logos generated by the SMT for the noisy and noiseless data sets exhibited a considerable similarity to the logos provided by JASPAR, with some small differences observed. This similarity indicates that the logos generated by the SMT can serve as initial seeds for optimization algorithms, such as the Expectation-Maximization (EM) algorithm, enabling further improvements in the quality of the logos. The consistency in the representation of the logos in the noisy and noiseless data sets also demonstrates the SMT’s ability to efficiently handle the presence of noise in the analyzed genomic data.

Through these experiments, we employed an algorithm called *n-seeds* that operates on the SMT and efficiently extracts all the most enriched k-mers and all k-mers with up to d mutations. This algorithm is particularly useful for efficiently constructing positional weight matrix (PWM) models, as it allows the identification of conserved regions and recurring patterns in DNA and RNA sequences.

The results obtained demonstrate that the SMT presents robust performance even in situations with noisy data, indicating its applicability in real-world scenarios where data quality can vary. The implementation of the n-seeds algorithm on the SMT highlights the versatility of the data structure in motif discovery and PWM model construction. This innovative approach has the potential to significantly improve the analysis of large-scale genomic data, contributing to advances in the field of bioinformatics and genomics.

In summary, the results obtained in this section demonstrate the effectiveness of SMT in handling ChIP-seq datasets, as well as its ability to adapt to noisy data. The application of the n-seeds algorithm on SMT further illustrates its potential as a powerful tool in motif discovery and PWM model construction, paving the way for future research and applications in the field of genomics.

## Conclusion

Additionally, the implementation of the n-seeds algorithm on the SMT highlights its versatility in motif discovery and PWM modeling, providing a powerful tool for large-scale genomic data analysis. The results obtained in this work demonstrate the potential of the SMT and its applications in the analysis of ChIP-seq data, opening the way for further research and applications in the genomics field.

In this paper, we discussed the advanced features of SMT, including constant-time exact search, fast identification of most abundant kmers, and linear-time approximate search for finding fragments with up to *d* mutations uniformly distributed along their bases. These features demonstrate the versatility and effectiveness of SMT, making it an important tool for researchers dealing with large-scale genomic data analysis.

Although the SMT is a promising data structure, it is important to recognize that there is still room for improvement and future advancements. As new technologies and approaches are developed, the SMT and similar data structures should evolve to keep up with the growing demand for efficient and effective solutions in the field of bioinformatics and genomics. Continuous collaboration between researchers and developers is essential to ensure that the SMT remains a useful and updated tool for the discovery of motifs and other applications in the analysis of genomic data.

## Acknowledgments

The authors thank the anonymous reviewers for their valuable suggestions. This work is supported in part by funds from the National Science Foundation (NSF: # 1636933 and # 1920920).

## References

J. Archbold and N. Johnson. A construction for room’s squares and an application in experimental design. The Annals of Mathematical Statistics, 29(1):219–225, 1958.

T. L. Bailey, J. Johnson, C. E. Grant, and W. S. Noble. The meme suite. Nucleic acids research, 43(W1):W39–W49, 2015.

J. A. Castro-Mondragon, R. Riudavets-Puig, I. Rauluseviciute, R. Berhanu Lemma, L. Turchi, R. Blanc-Mathieu, J. Lucas, P. Boddie, A. Khan, N. Manosalva Pérez, et al. Jaspar 2022: the 9th release of the open-access database of transcription factor binding profiles. Nucleic acids research, 50(D1): D165–D173, 2022.

M. J. Chaisson, A. D. Sanders, X. Zhao, A. Malhotra, D. Porubsky, T. Rausch, E. J. Gardner, O. L. Rodriguez, L. Guo, R. L. Collins, et al. Multi-platform discovery of haplotype-resolved structural variation in human genomes. Nature communications, 10(1):1784, 2019.

S. Deorowicz, A. Gudyś, M. Długosz, M. Kokot, and A. Danek. Kmer-db: instant evolutionary distance estimation. Bioinformatics, 35(1):133–136, 2019.

S. Goodwin, J. D. McPherson, and W. R. McCombie. Coming of age: ten years of next-generation sequencing technologies. Nature Reviews Genetics, 17(6):333–351, 2016.

K. L. Howe, P. Achuthan, J. Allen, J. Allen, J. Alvarez-Jarreta, M. R. Amode, I. M. Armean, A. G. Azov, R. Bennett, J. Bhai, et al. Ensembl 2021. Nucleic acids research, 49 (D1):D884–D891, 2021.

C. Marchet, L. Lecompte, C. D. Silva, C. Cruaud, J.-M. Aury, J. Nicolas, and P. Peterlongo. De novo clustering of long reads by gene from transcriptomics data. Nucleic Acids Research, 47(1):e2–e2, 2019.

C. Sanderson and R. Curtin. Armadillo: a template-based c++ library for linear algebra. Journal of Open Source Software, 1(2):26, 2016.

D. E. Wood, J. Lu, and B. Langmead. Improved metagenomic analysis with kraken 2. Genome biology, 20:1–13, 2019.

