## Supplemental Data for "Discovering motifs and genomic patterns with SMT: a high-performance data structure for counting kmers"

### Supplementary materials

Jader M. Caldonazzo Garbelini

**Abstract**

In this supplementary material, we present the algorithms that have been employed in the main paper. The algorithms showcased here include: create-SMT, which demonstrates how the SMT is constructed; k-SEARCH, which illustrates how the exact search is performed; d-DFS:, which explains how the approximate search is carried out; s-DLT: which demonstrates how kmers are removed from the SMT; and finally, the get-seeds, which returns the enriched kmers from the SMT. These algorithms are fundamental to the implementation and analysis of the methods described in the study. The purpose of this material is to provide a detailed insight into the algorithmic approaches used and to facilitate the understanding and reproduction of the results presented in the article.

### 1 Create SMT

Algorithm 1 describes the process of creating a Sparse Motif Tree (SMT) from a set of sequences. Using this algorithm, it is possible to build an efficient tree that represents the k-mers present in the sequences, optimizing the use of memory and computing time. The SMT is essential to enable exact and approximate searches in constant and linear time, respectively, facilitating the analysis of large-scale genomic data.

**Algorithm 1** *create-SMT*:

```
1:  for s ∈ S do
2:     node = root
3:     for i = 1 to k do
4:        symbol = s[i]
5:        if M[node, symbol] = 0 then
6:           M[node, symbol] = new_node
7:           new_node = new_node + 1
8:           node = new_node
9:        else
10:          node = M[node, symbol]
11:       end if
12:    end for
13: end for
14: return M
```

The *create-SMT* algorithm builds an SMT from a set of sequences $S$. For each sequence $s$ in $S$ (Line 1), the algorithm starts with the root node of the tree (Line 2). Then, it traverses the symbols of sequence $s$, one by one, from the first to the k-th symbol (Line 3). For each symbol of the sequence (Line 4), the algorithm checks if there is a corresponding child node for this symbol in the matrix $M$ (Line 5). If there is no corresponding child node (i.e., $M[node, symbol] = 0$), the algorithm creates a new node (Line 6), updates the matrix $M$ by associating the new node with the current symbol (Line 7), increments the index of the new node (Line 8), and sets the current node to the new node created (Line 9). If there already exists a corresponding child node, the algorithm simply updates the current node to that child node (Line 10-11). The process is repeated for all the sequences in $S$ and all the symbols in the sequences. At the end of the execution, the algorithm returns the matrix $M$ (Line 13), which represents the structure of the SMT built from the set of sequences $S$.

### 2 Search on SMT

Algorithm 2 describes the process of exact search in the SMT, detailing how a specific k-mer is located within the data structure. This algorithm allows for identifying the presence or absence of a k-mer in the SMT, providing crucial information about patterns and conserved regions in the analyzed sequences.

**Algorithm 2** k-SEARCH

```
1: for i = 1 to k do
2:     symbol = kmer[i]
3:     next = SMT(symbol, node)
4:     if next ≠ 0 then
5:         node = next
6:     else
7:         return False
8:     end if
9: end for
10: return True
```

The k-SEARCH algorithm details the process of exact search for a k-mer within the SMT. Initially, the algorithm traverses each symbol of the k-mer, from index 1 to k (line 1). For each symbol, the algorithm searches for the corresponding node in the SMT (line 2) and checks if the next node exists (line 3). If the next node exists, the algorithm moves to the next node (line 4). Otherwise, the search fails, and the algorithm returns "False" (line 5). If all symbols of the k-mer are found in the SMT, the search is successful, and the algorithm returns "True" (line 8).

Algorithm 3 describes the process of approximate search in the Sparse Motif Tree (SMT). Using a depth-first search approach with error tolerance (d-DFS), the algorithm explores the paths in the SMT, allowing for a limited number of errors along the path. This enables the identification of sequences that approximately match the searched k-mer, even when there are variations or mutations.

**Algorithm 3** *d-DFS*$(M, s, d\_max, d, i, k, node, resp)$:

```
1: if d > d_max : then
2:     return    // The search string exceeded d mismatches and the tree will be pruned.
3: end if
4: if i ≥ k : then
5:     resp = 1    // The search string was found with up to d mismatches.
6: end if
7: symbol = s[i]
8: for j = 1 to 4 do
9:     next_node = M[node, j]
10:    if next_node ≠ 0 and symbol = j then
11:        d-DFS(M, s, d_max, d, i + 1, k, next_node, resp)    // Match!
12:    else
13:        d-DFS(M, s, d_max, d + 1, i + 1, k, next_node, resp)    // Mismatch!
14:    end if
15:    if resp = 1 then
16:        break
17:    end if
18: end for
```

The d-DFS algorithm starts at line 1 by checking if the number of allowed errors ($d$) has exceeded the maximum error limit (d_max). If this happens, the search is interrupted and the tree is pruned. If the character position ($i$) is greater than or equal to the k-mer size ($k$), the search was successful, and the function returns successfully (lines 3-5).

At line 6, the algorithm extracts the current symbol from the search sequence ($s$). The algorithm then iterates over the possible symbols ($j$) in lines 7-18. For each symbol, the algorithm checks if the next node in the tree exists (line 9). If the next node exists and the current symbol matches the symbol in the search sequence, the algorithm proceeds with the recursive search, maintaining the same error count ($d$) (line 10). Otherwise, the algorithm proceeds with the recursive search, increasing the error count by 1 ($d + 1$) (line 12). The search ends when a successful path is found, i.e., when the variable $resp$ is set to 1 (lines 14-16).

### 3 Update SMT

Algorithm 4 describes the process of removing k-mers and their surroundings in the SMT. This algorithm is essential for refining and cleaning the SMT, ensuring that only relevant and conserved k-mers are retained in the data structure. The algorithm efficiently removes k-mers and allows for subsequent analysis of conserved regions and recurring patterns in DNA and RNA sequences.

**Algorithm 4** s-DLT($SMT, node, kmer, j, k$)

```
1:  if j ≥ k then
2:     SMT[node, $] = 0    // Remove fragment by setting its value to 0.
3:  else
4:     symbol = kmer[j]
5:     if symbol == X then
6:        for symbol = 1 to 4 do
7:           next = SMT[node, symbol]
8:           if next ≠ 0 then
9:              s-DLT(SMT, next, kmer, j + 1, k)
10:          end if
11:       end for
12:    else
13:       next = SMT[node, symbol]
14:       if next ≠ 0 then
15:          s-DLT(SMT, next, kmer, j + 1, k)
16:       end if
17:    end if
18: end if
```

The *s-DLT* algorithm starts at line 1, checking if the current index $j$ is greater than or equal to the k-mer size $k$. If true, line 3 removes the fragment by setting its value to 0. Otherwise, the algorithm continues at line 6, where the current symbol of the k-mer is assigned to the variable *symbol*. If the current symbol is equal to $X$, indicating a wildcard position, the algorithm enters a loop at line 8 for all four possible symbols. For each symbol, it checks at line 10 if the next node in the SMT is not equal to 0. If it is not, the s-DLT function is called recursively at line 11 for the next node, k-mer, and incremented index. If the current symbol is not $X$, the algorithm checks at line 14 if the next node in the SMT is not equal to 0. If it is not, the *s-DLT* function is called recursively at line 15 for the next node, k-mer, and incremented index. This process continues until all specified fragments are removed from the SMT.

### 4 Build models

Algorithm 5 describes the process of extracting enriched k-mers from the SMT. It details the method used to identify and collect significant k-mers based on a set of statistical criteria, as well as removing surrounding k-mers from the SMT. This algorithm is crucial for extracting useful and relevant information from analyzed DNA and RNA sequences, enabling researchers to gain a deeper understanding of the conserved properties of biological sequences and facilitating large-scale analysis of genomic datasets.

**Algorithm 5** get-SEEDS

```
1: while j < nseeds do
2:   result = getBestKmer(SMT, fasta, k)
3:   kmer = result[kmer]
4:   obs = result[count]
5:   p = getBKprob(β, kmer)
6:   exp = p * m * n
7:   pval = fisher_test(obs, exp, mn − obs, mn − exp)
8:   if pval > sig_level then
9:     break
10:  end if
11:  seeds[kmer] = obs
12:  surround(kmer, x)
13:  surroundDelete(SMT, kmer, k)
14:  ++j
15: end while
16: return seeds
```

The *get-SEEDS* algorithm initially establishes a loop that continues until the desired number of seeds (*nseeds*) is reached (line 1). During each iteration of the loop, the algorithm first obtains the best k-mer and its observed count (lines 2 and 3), as well as the base probability ($p$) and expected count (*exp*) (lines 4 and 5). The algorithm then calculates the p-value using the Fisher's exact test (line 6). If the p-value is greater than the established significance level (line 7), the loop is terminated (line 8). Otherwise, the k-mer is added to the seeds (line 9) and the surrounding k-mers are removed from the SMT (lines 10 and 11). The counter $j$ is incremented (line 12) and the loop continues until all desired seeds are obtained. Finally, the algorithm returns the found seeds (line 14).

Algorithm 6 describes the process of creating models from the k-mers extracted from the SMT. This process is carried out by constructing PWM matrices for each found seed, and the algorithm details how these matrices are generated and stored from the k-mers and their corresponding counts. At the end of the process, a list of constructed models is returned, allowing for analysis and identification of conserved patterns in genomic sequences.

**Algorithm 6** bs-MODELS

```
1: for seed ∈ seeds do
2:   siblings = d-DFS (SMT, seed, d)
3:   kmers = siblings[kmers]
4:   counts = siblings[counts]
5:   for i = 1 to |kmers| do
6:     kmer = kmers[i]
7:     count = counts[i]
8:     for j = 1 to k do
9:       symbol = kmer[j]
10:      PCM[symbol, j] += count
11:    end for
12:    PWM = get_pwm(PCM)
13:    models.append(PWM)
14:  end for
15: end for
16: return models
```

In the *bs-MODELS* algorithm, for each *seed* in *seeds* (line 1), an approximate search (*d-DFS*) is performed with the SMT, the seed, and a maximum Hamming distance limit $d$ (line 2). The result includes the found k-mers (*siblings[kmers]*) and their corresponding counts (*siblings[counts]*) (lines 3 and 4). Then, for each found k-mer (line 5), the algorithm updates the position count matrix (PCM) according to the k-mer count and its position in the sequence (lines 6-10). The PCM matrix is then converted into a position weight matrix (PWM) (line 11), and the resulting PWM model is added to the model list (line 12). The algorithm returns the constructed model list (line 15).

### 5 Conclusion

The SMT and its associated algorithms provide an efficient and robust solution for the analysis of large volumes of genomic data. The SMT enables rapid and accurate identification of k-mers, as well as detection of conserved patterns and analysis of variations within DNA and RNA sequences. With the increasing demand for tools capable of handling the exponential growth of genomic data, the SMT and its satellite algorithms prove to be a valuable alternative for researchers and bioinformatics professionals, contributing to a better understanding of gene functions and regulations and aiding in the identification of functional and structural elements of the genome.
